# Polycomb Repressive Complex 1.1 Component, BCOR, Promotes Syncytiotrophoblast Differentiation in Mice and Humans

**DOI:** 10.1101/2024.01.29.577740

**Authors:** Danielle Sadowski, Connie M. Corcoran, Riyan Abdi, Teng Zheng, Hiroaki Okae, Takahiro Arima, Vivian J. Bardwell, Micah D. Gearhart

## Abstract

Early defects in placenta development are thought to underlie a range of adverse pregnancy conditions including miscarriage, fetal growth abnormalities, preeclampsia, and stillbirth. Differentiating trophoblast stem cells undergo a choreographed allocation of syncytiotrophoblast and extravillous trophoblast cells in response to signaling cues from the developing fetus and the uterine environment. The expression and activity of transcription factors and chromatin modifying enzymes change during differentiation to appropriately reshape the chromatin landscape in each cell type. We have previously found in mice that extraembryonic loss of BCOR, a conserved component of the epigenetic silencing complex Polycomb Repressive Complex 1.1 (PRC1.1), leads to a reduced labyrinth and expanded trophoblast giant cell population in the placenta. Molecular analysis of wild-type and BCOR loss-of-function male and female placentas by RNA-seq identified gene expression changes as early as E6.5. We found that BCOR is required to down regulate stem cell genes and repress factors that promote alternate lineages which leads to reduced levels of syncytiotrophoblasts. ChIP-seq experiments identified a number of directly bound functional targets including *Pdgfa* and *Wnt7b*. In humans, *BCOR* is mutated in X-linked syndromes involving fetal growth restriction and females with a heterozygous null mutation in *BCOR* can experience recurrent miscarriages. To establish a direct role for *BCOR* in human placental development, we used CRISPR/Cas9 to knockout *BCOR* in male (CT29) and female (CT30) human trophoblast stem cells. Mutant cell lines retained capacity for induced differentiation into syncytiotrophoblast and extravillous trophoblasts and exhibited minimal changes in gene expression. However, in 3D cell culture using trophoblast organoid media, *BCOR* knockout lines had significantly altered gene expression including homologs of stem cell genes upregulated in *Bcor* knockout mice. CUT&RUN experiments in self-renewing and 3D cell culture identified genes directly bound by BCOR. Single cell profiling of wild type, knockout, and a P85L pathogenic knock-in *BCOR* mutation showed a reduced capacity to differentiate into syncytiotrophoblasts after four days of differentiation. Together, these results suggest that BCOR is a conserved regulator of trophoblast development that represses stem cell genes during differentiation and maintains lineage fidelity by repressing genes that promote alternate cell fates.

## Introduction

The trophectoderm gives rise to the main extraembryonic structures of placenta as well as components of the parietal yolk sac and chorion. Various cells within the placenta are responsible for pregnancy hormone production and the remodeling of maternal arteries facilitating the transport of oxygen and nutrients and the removal of waste products to support fetal development. Faithful allocation of these cell types is required for successful placentation and, conversely, inappropriate differentiation can lead to pregnancy complications including fetal growth restriction, preeclampsia and increased maternal and fetal mortality. Placental insufficiencies can also limit nutrient and growth support for the fetus which can lead to low birth weights, chronic neurodevelopmental disorders(Bronson and Bale 2016), and congenital heart defects(Pan and Li 2023). Few molecular factors that control trophoblast differentiation are well understood, limiting the development of diagnostic and therapeutic strategies to reduce complications that occur during pregnancy and postnatal development.

BCOR is a core component of Polycomb Repressive Complex 1.1(Gearhart et al. 2006; Gao et al. 2012). The PRC1 family of complexes have a conserved evolutionary role in maintaining lineage identity by modifying chromatin structure (Bracken and Helin 2009; Kim and Kingston 2022). Although somatic *BCOR* loss-of-functions mutations are frequently found in various types of cancer (reviewed in (Astolfi et al. 2019)), germline mutations are much more rare, occurring less than once in a million live births (“MedlinePlus” 2024). These mutations cause a range of syndromes characterized by microphthalmia, cognitive delay and craniofacial and heart defects(Ng et al. 2004; Ragge et al. 2019). Because *BCOR* is located on the X-chromosome, females with Oculofacialcardiodental syndrome (OFCD) exhibit a mosaic expression of wild type and mutant alleles depending on random X-inactivation. Skewing (90-100%) of leukocytes expressing the wild type *BCOR* allele has been observed in OFCD patients(Ng et al. 2004) and presumably also occurs in their placentas. These patients can experience fetal growth restriction at birth and recurrent miscarriages as adults which are likely due to embryonic lethality of male offspring(Ragge et al. 2019). Male patients, including those with Lenz microphthalmia, most often carry hemizygous missense mutations and can have neurocognitive delay and fetal growth restriction(Ng et al. 2004; Lenz 1955; Kraus et al. 2018; X. Zhu et al. 2015; Du et al. 2018). For example, a recently reported male patient born during the 39th week of gestation and weighing 1,860g (-3.7 SD) was found to have a mutation introducing a stop codon in the last exon(Schwaibold, Brugger, and Wagner 2021).

In contrast to humans, the paternal X-chromosome is selectively inactivated in extraembryonic tissues of rodents(Takagi and Sasaki 1975; West et al. 1977). Breeding *Bcor* ^Flox910^ mice so that extraembryonic tissues expressed a wild type allele produced viable *Bcor* ^Δ910/WT^ pups (Hamline et al. 2020), suggesting that placental insufficiencies are responsible for prenatal lethalities in female mice expressing the mutant *Bcor* from the maternal allele in extraembryonic tissues. Previous studies have shown that loss of *Bcor* in cultured mouse trophoblast stem cells causes upregulation of *Eomes* expression and promotes syncytiotrophoblast differentiation in an activin dependent manner (G. Zhu et al. 2015). We previously reported that mutations in X-linked *Bcor* gene lead to placental insufficiency in mice on a mixed background (Hamline et al. 2020). To determine the molecular nature of these defects here we profile characterize the localization and genome wide occupancy of BCOR as well as gene expression changes upon extraembryonic loss of the maternal *Bcor* allele in mice. We also model loss-of-function and missense mutations in *BCOR* using human trophoblast stem cells. Together these results show that BCOR is an important regulator of trophoblast differentiation in mice and humans.

## Results

We performed immunofluorescence staining to identify which cell types express BCOR in the C57BL/6 mouse placenta. In the earliest time point examined, E6.5, we see strong BCOR expression in the extraembryonic ectoderm, the ectoplacental cone and parietal trophoblast giant cells (**Fig 1A**). At E7.5 BCOR expression persists in the ectoplacental cone and parietal trophoblast but can also be seen in the developing chorion. Notably, BCOR staining is absent from the hinge regions characterized by the expression of high levels of the stem cell markers *Eomes* and *Sox21(Kuales et al. 2015)*. By E8.5, BCOR expression is highest in the chorionic ectoderm although expression is still observed in a subset of parietal trophoblast giant cells. This protein expression is consistent with the mRNA expression we have reported at E7.5 and E8.5 on a mixed background(Wamstad and Bardwell 2007; Hamline et al. 2020).

**Figure 1.**
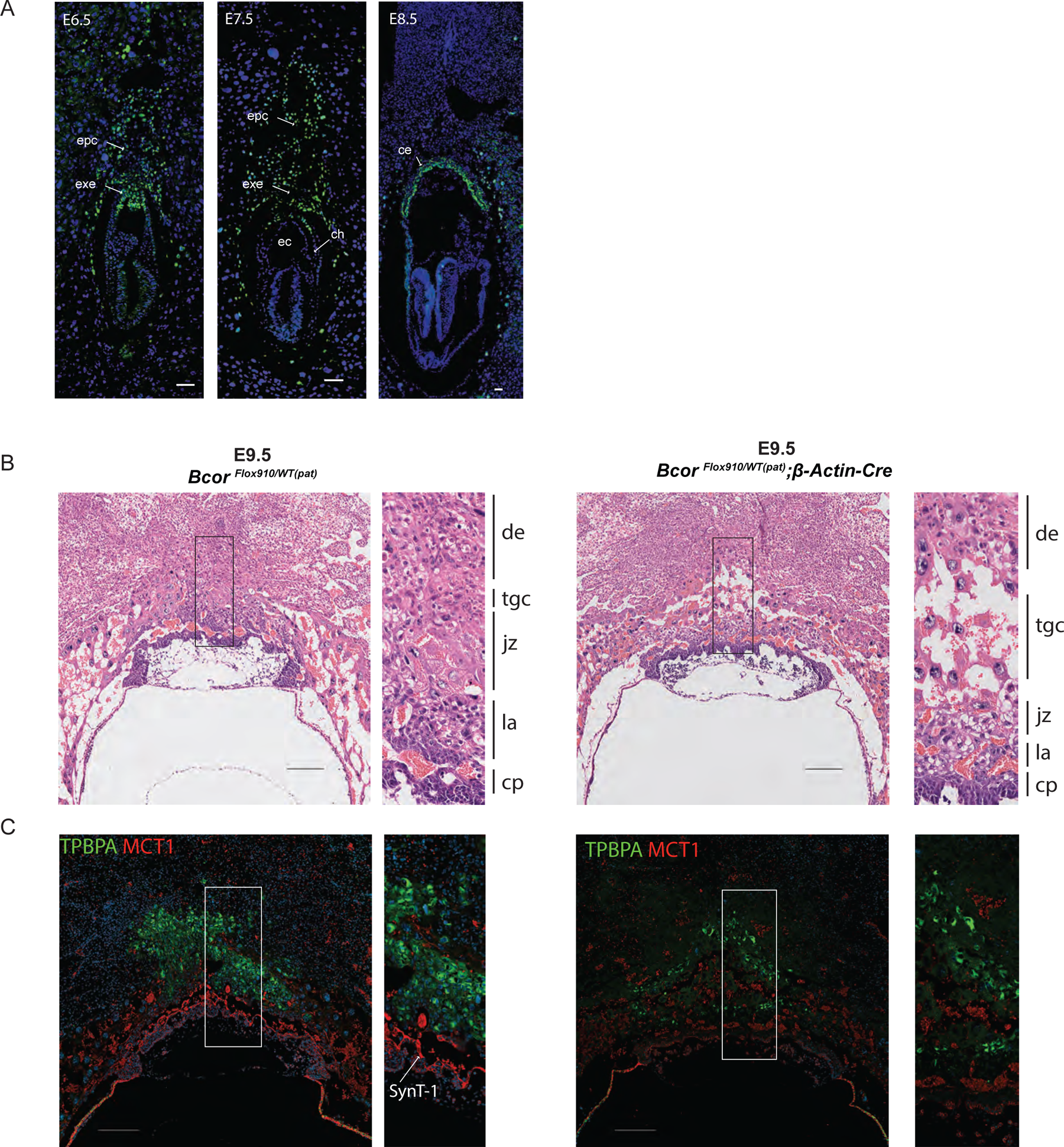
BCOR Expression in the Mouse Placenta. A) Immunofluorescence of BCOR expression (green) and DAPI (blue) of cryosectioned concepti from E6.5 to E8.5. BCOR is strongly expressed in the nuclei of cells in the ectoplacental cone (epc) and extraembryonic ectoderm (exe) at the early time points. At E7.5 BCOR expression is present in the developing chorion but absent from the chorionic hinge (ch). BCOR is expressed throughout the chorionic ectoderm (ce) at E8.5. The exocoelomic cavity (ec) is labeled at E7.5. Scale bars (white) are 50 μm in each panel. B) Hematoxylin and eosin staining of medial sections from paraffin embedded *Bcor* ^Flox910/WT(pat)^ (left) and *Bcor* ^Flox910/WT(pat)^;β-Actin-Cre (right) concepti at E9.5. Expanded regions shown to the right of each panel are labeled with the locations of chorionic plate (cp), labyrinth (la), junctional zone (jz), trophoblast giant cells (tgc), and decidua (de). Scale bars (black) are 200 μm in each panel. C) MCT1 (red) and TPBPA (green) staining of *Bcor* ^Flox910/WT(pat)^ (left) and *Bcor* ^Flox910/WT(pat)^;β-Actin-Cre (right) concepti at E9.5. Background cytoplasmic staining is present in erythrocytes on the red channel. Scale bars are 200 μm in both panels.

We mated homozygous *Bcor* ^Flox910/Flox910^ females that had been backcrossed onto a C57BL/6 background with C57BL/6 males expressing Cre recombinase driven by the β-*Actin* promoter (β-*Actin*-Cre). Because the paternal X chromosome carrying a wild type copy of *Bcor* will be inactivated in extraembryonic cells, the maternal *Bcor* ^Flox910^ allele will be the only one expressed throughout most of the placenta. We previously found that the paternal *Bcor* gene is able to escape X-inactivation in parietal trophoblast giant cells in the ectoplacental cone at E8.5 in a mixed 129 background (Hamline et al. 2020). Hemizygous male pups, however, only carry the maternal floxed allele (*Bcor* ^Flox910/Y^) and in pups expressing β-*Actin*-Cre will have complete loss of full length *Bcor* in the placenta as well as the embryo at time points prior to fetal demise at ∼E8.5. Hematoxylin and eosin (H&E) staining of median sagittal sections *in utero* confirmed that previous observations of a reduced labyrinth layer and increased numbers of trophoblast giant cells in *Bcor* ^Flox910/WT^;β-*Actin*-Cre females at E10.5 (Hamline et al. 2020) are also present at E9.5 in this inbred background (**Fig. 1B**). To examine which cell types are under or overrepresented in the mutants we stained E9.5 sections with MCT1 and TPBPA to label SynT-I and subsets of trophoblast giant cells, respectively (Nagai et al. 2010; Simmons, Fortier, and Cross 2007). We find that both MCT1 and TPBPA are expressed in fewer cells in *Bcor* ^Flox910/WT^;β-*Actin*-Cre females compared to control animals (**Fig. 1C**). This suggests that the reduced labyrinth size could be due to a compromised SynT-I layer and the expansion of trophoblast giant cells is likely derived from a TPBPA negative population such as primary parietal trophoblast giant cells(Simmons, Fortier, and Cross 2007).

To determine which genes are regulated by BCOR that could be contributing underdeveloped labyrinth and expanded giant cell populations, we performed bulk RNA-seq on individual placenta at E6.5 and E7.5 for Bcor ^Flox910/Y^;β-actin-Cre and Bcor ^Fl/Y^ males and E6.5-E9.5 *Bcor* ^Flox910/WT^;β-actin-Cre and *Bcor* ^Flox910/WT^ females prior to the fetal demise at E8.5 and E12.5 in each sex respectively. The samples contained polar extraembryonic tissue at each time point, i.e. ectoplacental cone at E6.5 and chorion and extraembryonic ectoderm at E8.5-E9.5. Four replicates were prepared from individual mice at each timepoint from each group although five low quality samples were omitted leaving a minimum of three replicates in each group. Analysis of genes previously characterized in the mouse placenta provided temporal framing for our dataset. For example, *Brachyury* (*T*) was detected in samples collected at E7.5 suggesting extraembryonic mesoderm was present in our dissections. Similarly, *Vcam1* was present in samples collected at E8.5 and E9.5 reflecting chorioallantoic fusion had occurred consistent with previously reported timing(Gurtner et al. 1995). We next used principal component analysis of female and male placentas to evaluate the differentiation of wild type and *Bcor* knockout placenta samples (**Fig. 2A**). The first principal component, PC1, captured the variance across time for both sexes and showed that while samples from each time point clustered separately, the knockout placentas lagged behind wild type samples starting at E7.5. For example, wild type female E8.5 samples (red diamonds) were all closer to the E9.5 cluster than the E8.5 knockout samples (blue diamonds). Similarly, the wild type E7.5 males (red triangles) are further away from the E6.5 samples than the Bcor^Null/Y^ knockout E7.5 samples (blue triangles). These findings suggest *Bcor* mutant placentas have a slight delay but remain most similar to wild type placentas collected at the same time point.

**FIgure 2.**
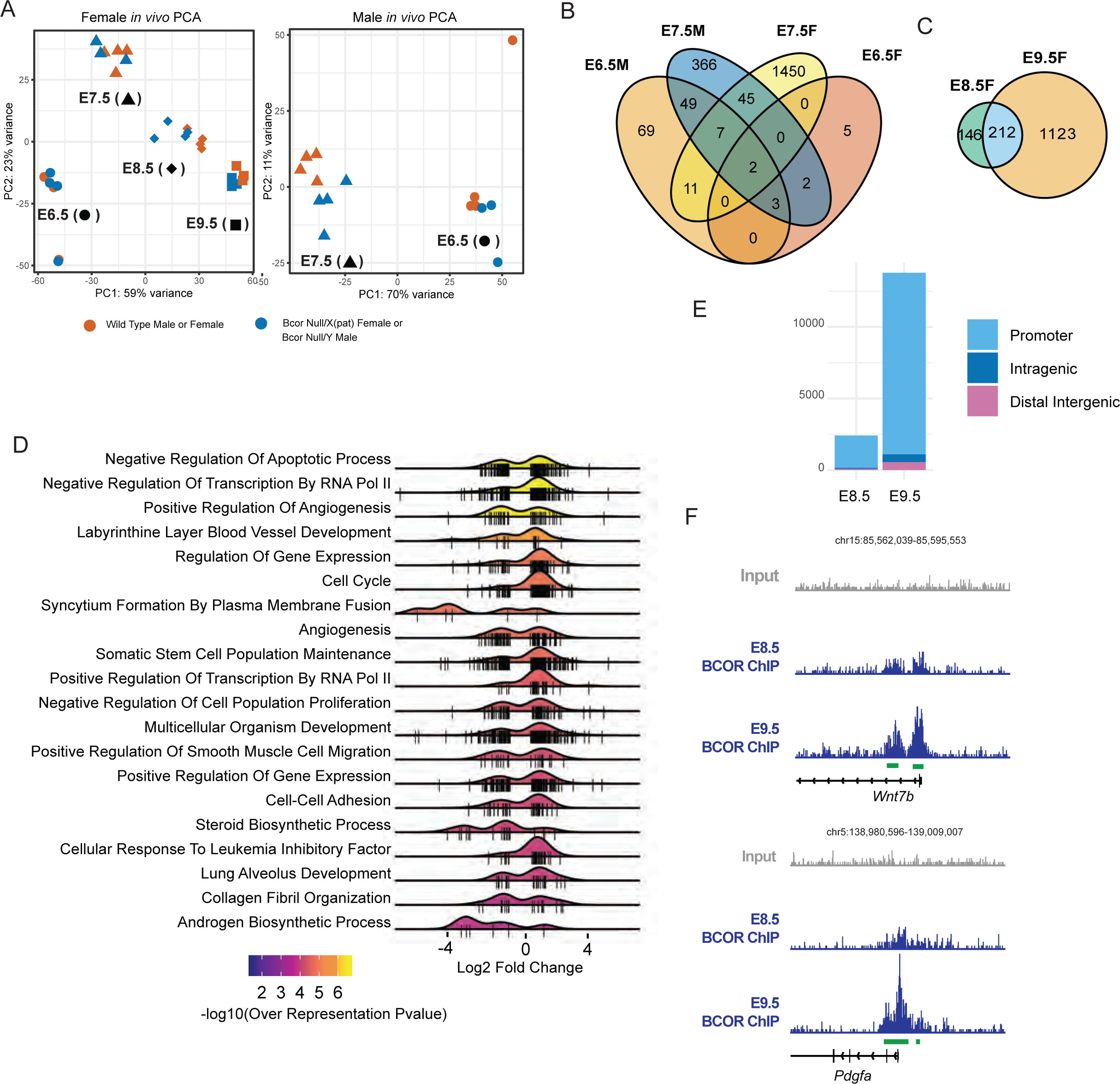
Genome Wide Profiling of Mouse Placenta. A) Principal component analysis gene expression data from individual female (left) and male (right) placentas. Control animals are shown in red and mutant animals (*Bcor* ^Flox910/WT(pat)^;β-Actin-Cre or *Bcor* ^Flox910/Y^;β-Actin-Cre) are shown in blue. Embryonic days post-coitum are indicated with ⏺ (E6.5), ▲(E7.5), ⬥(E8.5), or (E9.5). B) Venn diagram of overlapping genes at E6.5 and E7.5 for female and male placentas. C) Venn diagram of overlapping genes at E8.5 and E9.5 for female placentas. D) Top 20 enriched gene ontology terms for genes differentially expressed in one or more sample groups. The ridgeline plots on the right show the distribution of log_2_ fold changes for the overrepresented genes in each term. If a gene was differentially expressed in more than one sample, the log_2_ fold change from sample with most significant *p* values was chosen. The colors under the ridgeline plot correspond to the -log_10_(*p* value) for the over representation of each particular gene ontology term. E) Distribution of BCOR peaks at E8.5 and E9.5. Promoter associated peaks were defined as any peak within 3kb upstream or downstream of the transcription start site. F) Examples of BCOR ChIP-seq tracks at the promoters *Wnt7b* and *Pdgfa*.

We next looked at *Bcor* expression across time. In wild type males, the expression was 35±12 and 44±6 FPKM at E6.5 and E7.5 respectively. In wild type females, Bcor expression changed from 59±22 to 51±6, 40±4, and 18±1 FPKM as development progressed from E6.5 to E9.5. The high variation at E6.5 likely reflects inconsistencies in the dissections due to the small size. However, the downward trend is consistent with immunofluorescence experiments, reflecting the fraction of BCOR+ cells present in the samples. The floxed exons, 9 and 10, contain 584 nucleotides in total which is 8.4% of the 6,941 nucleotide *Bcor* transcript. This would presumably not have a significant effect on the total number of reads mapping to the transcript. Yet for the male knockout samples, *Bcor* was expressed at much lower level: 11±1 and 5±1 FPKM at E6.5 and E7.5 respectively, suggesting the premature stop codon introduced by the loss of these exons leads to nonsense mediated decay. In female knockouts the change in *Bcor* expression was less pronounced, changing from 17±5 to 27±12, 16±6 and 14±2 FPKM from E6.5 to E9.5 respectively. These more subtle changes likely reflects the contribution from extraembryonic mesoderm at E7.5 or allantois at E8.5 and E9.5 derived from the embryo proper expressing *Bcor* mosaically from either allele and/or a contribution due to escape from X-inactivation of the paternal allele in parietal-trophoblast giant cells as has been observed previously(Hamline et al. 2020).

Given the differences between male and female samples, we determined differentially expressed genes for each time point independently for each sex. Using a cutoff of 1.5-fold fold change and a Benjamini-Hochberg adjusted *p* value of 0.05, we found 12, 1515, 358, and 1335 differentially expressed genes in female placentas at E6.5, E7.5, E8.5, and E9.5 respectively. For male concepti, 141 and 474 genes were differentially expressed at E6.5 and E7.5 respectively. For E6.5 and E7.5, we sought to identify genes that were differentially expressed in both sexes. We identified 5 genes at E6.5 and 54 genes at E7.5, both including *Bcor* itself, indicating that these timepoints represent very early downstream consequences of BCOR loss (**Fig. 2B**). For example, *Eomes* is upregulated by 4.6-fold (*p*_adj_=2.5^e-3^) and 3.5-fold (*p*_adj_=9.4^e-8^) at E6.5 in males and females respectively. This gene was previously found to be repressed by BCOR in shRNA experiments using mouse trophoblast stem cells(G. Zhu et al. 2015). After allantoic attachment in female samples, we identified 212 differentially expressed genes present at both timepoints (**Fig. 2C**). These genes include *Tfap2a*, up 3-fold (*p*_adj_=4.3^e-6^) and 4.3-fold (*p*_adj_=5.4^e-34^), and *Wnt7b*, up 1.8-fold (*p*_adj_=1.3^e-5^) and 3.8-fold (*p*_adj_=3.3^e-30^), at E8.5 and E9.5 respectively which have previously been shown to regulate trophoblast differentiation (Krendl et al. 2017; Parr et al. 2001).

To understand which pathways are regulated by BCOR we tallied 3,125 genes differentially expressed at one or more time points to identify statistically overrepresented gene ontology (GO) terms describing biological processes (**Fig. 2D**). We found that regulation of apoptosis, transcriptional regulation and angiogenesis among the top GO terms. Notably, *labyrinth layer blood vessel development* (GO:0060716, p=3.1^e-6^) and *syncytium formation by plasma membrane fusion* (GO:0000768, *p*=1.4^e-5^) were highly enriched. These terms included a mixture of upregulated and downregulated genes (**Fig. 2D**, right). *Synb* and *Gcm1* were found to be downregulated at E9.5 (1.6-fold down, *p*_adj_=9.7^e-6^ and 1.6-fold down, *p*_adj_=1.4^e-2^, respectively) in contrast to *in vitro* shRNA experiments showing these genes were upregulated(G. Zhu et al. 2015). These decreases likely reflect a proportional reduction of cells in labyrinth layer within the dissected sample while stem cell markers, such as *Satb1(Yu et al. 2022)* (1.6-fold up, *p*_adj_=1.2^e-6^) and *Tet1(Senner et al. 2020)* (1.9-fold up, *p*_adj_=1.9^e-6^), are proportionally increased.

To determine which of the upregulated genes were directly bound by BCOR we performed ChIP-seq from whole wild type placentas pooled over multiple litters and both sexes. We found 2,412 and 13,790 peaks at E8.5 and E9.5 respectively. As observed in hESC (Wang et al. 2018), the majority of the peaks were located at gene promoters (**Fig. 2E**). Of 2,025 genes upregulated in our dataset, 1,201 (59%) were directly bound by BCOR at E8.5 or E9.5. Among these functional targets, *Wnt7b* and *Pdgfa* were bound by prominent BCOR peaks at their respective transcriptional start sites at both time points (**Fig. 2F**). This data suggest that BCOR controls trophoblast development in mouse placenta by directly repressing key genes in trophoblast development.

Because mutations in *BCOR* that are found in OFCD are associated with recurrent miscarriages and intrauterine growth restriction (Ragge et al. 2019) we hypothesized that BCOR may also be required for trophoblast differentiation in humans. To test this hypothesis, we generated loss of function and missense mutations in male (CT29) and female (CT30) human trophoblast stem (hTS) cells (Okae et al. 2018) using CRISPR/Cas9. For BCOR knockouts, we used three control guide RNAs and three independent guide RNAs targeting exon six of the BCOR gene to generate hemizygous (CT29) and homozygous (CT30) loss of function mutations in the BCOR gene. Single colonies were isolated by limited dilution and screened by western blotting. Because the epitope of our in-house BCOR antibody recognizes residues in exons seven through nine, frame-shift mutations in exon six produce proteins lacking this epitope (**Fig. 3A**). Loss-of-function BCOR mutations were confirmed by Sanger sequencing.

**Figure 3.**
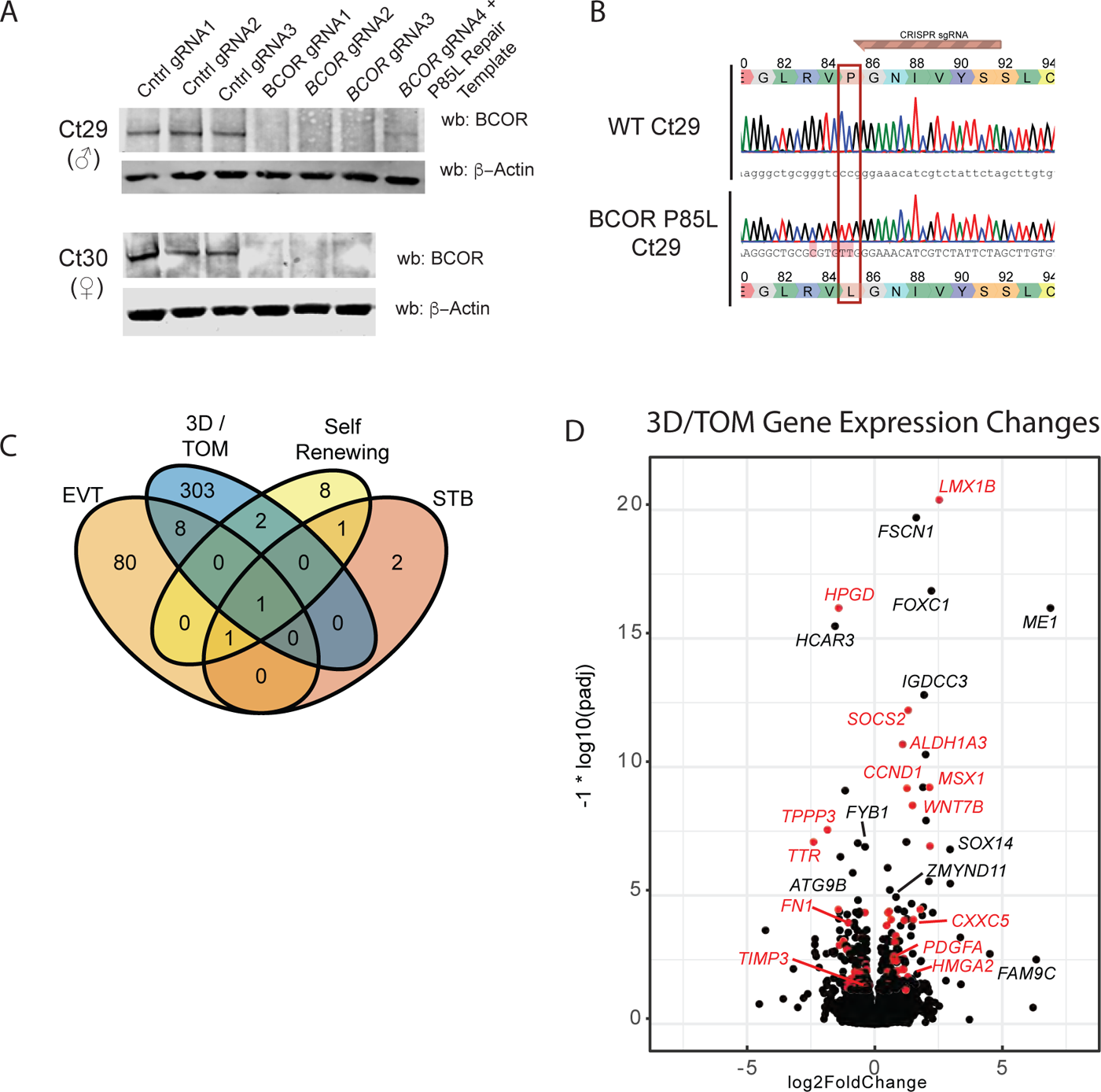
CRISPR/Cas9 Editing and Gene Expression of human trophoblast stem cells. A) Western blots of individual cell lines used in this study. Three independent controls and three guide RNAs targeting exon 6 of *BCOR* were used in CT29 (male) and CT30 (female cell lines). An additional guideRNA and single stranded repair template were used to engineer a proline to leucine mutation at amino acid 85 in the male cell line. The wobble positions of R83 and V84 were also mutated to facilitate detection by PCR. Sanger sequencing of genomic DNA (B) was used to confirm the mutation. The location of the gRNA protospacer used in this cell line is shown in brown. C) Four-way Venn diagram of differentially expressed genes in self renewing cells, extravillous trophoblasts (EVT), syncytiotrophoblast (STB) and autonomously differentiation cells in 3D culture using trophoblast organoid media (3D / TOM). D) Volcano plot of differentially expressed genes in 3D/TOM conditions. Log_2_FoldChange and *p* values were calculated using a mixed linear model with male and female samples incorporating sex as an independent variable. Genes shown in red were also differentially expressed at one or more timepoints in the *Bcor* ^Flox910^ mouse model.

In addition to BCOR loss-of-function mutations observed in females with OFCD, a reoccurring missense mutation at P85L has been observed in males(Ng et al. 2004; Ragge et al. 2019). To generate the P85L mutation we co-electroporated CT29 cells with synthetic single-stranded oligonucleotide 50 bases of homology on either side of the codon encoding proline 85. Additional silent mutations in the wobble positions of R83 and V84 were included so that clonal lines derived from single colonies could be screened by PCR. The mutation was confirmed by Sanger sequencing (**Fig. 3B**). Western blotting of this cell line side-by-side with wild-type and knockout cell lines indicates the P85L mutation affects BCOR stability and likely represents a strong hypomorphic allele.

Next we performed directed differentiation of 2D control and *BCOR* knock-out cells into syncytiotrophoblast and extravillous trophoblast using cAMP agonist and neureregulin-1 respectively(Strauss et al. 1992; Fock et al. 2015). We found that knock-out cells were competent for differentiation and were morphologically indistinguishable from control cell lines. We performed bulk RNA-seq on self-renewing cells, syncytiotrophoblast, and extravillous trophoblast and identified 13, 5 and 90 differentially expressed genes with a fold-change > 2 and an adjusted *p* value < 0.05 respectively, consistent with the observable morphological similarities. In contrast, seeding self-renewing control and *BCOR* knockout hTS cells in Matrigel for four days using media formulated for primary trophoblast organoids (TOM) (Turco et al. 2018) allowing for autonomous differentiation, led to a higher number of differentially expressed genes. Four days of culture was sufficient to capture the early stages of differentiation and the production in both control and knockout cell lines. *TEAD4* was downregulated 1.6 fold (*p*_adj_=1.8^e-13^) and syncytiotrophoblast genes such as *OVOL1* and *CGA* were upregulated 15.4 fold (*p*_adj_=1.9^e-60^) and 554 fold (*p*_adj_=4.8^e-192^) respectively in control cell lines. Comparing controls to BCOR knock-outs in CT29 and CT30 cell lines on day four we identified 314 differentially expressed genes including *LMX1B* (5.8-fold up, *p*_adj_=4.0^e-21^), *HMGA2* (2.2-fold up, *p*_adj_=7.5^e-3^), and *CXXC5* (2.9-fold up, *p*_adj_=8.6^e-5^) (**Fig. 3C**). Of these 314 human genes, 77 were also differentially expressed in the mouse (red dots, **Fig. 3D**), indicated that some of the pathways regulated by BCOR are conserved between species and that autonomous differentiation of hTS cells, within the context of 3D organoid-like cell culture, can recapitulate differentiation defects that occur *in vivo*.

To determine which genes were bound by BCOR in hTS cells, we performed CUT&RUN on BCOR using cells growing in 2D under self-renewing conditions. We found 1,722 BCOR peaks in self-renewing cells that were not observed in a knock-out cell line lacking the antibody epitope. To determine if the recruitment of other components of the PRC1.1 complex are dependent on BCOR, we also profiled KDM2B occupancy in wild-type and knockout cells. We identified 1,314 KDM2B peaks that co-occupy the 1,722 sites bound by BCOR. Of these 1,314 sites 1,251 sites remain bound by KDM2B in the absence of BCOR, including genes such as *SOCS2* which have a significant increase in gene expression in 2D self-renewing cells (2.2 fold, *p*_adj_=7.0^e-9^,**Fig. 4A**). This suggests that KDM2B, which contains a non-methylated CpG DNA-binding CXXC domain, is recruited to chromatin independently of BCOR.

**Figure 4.**
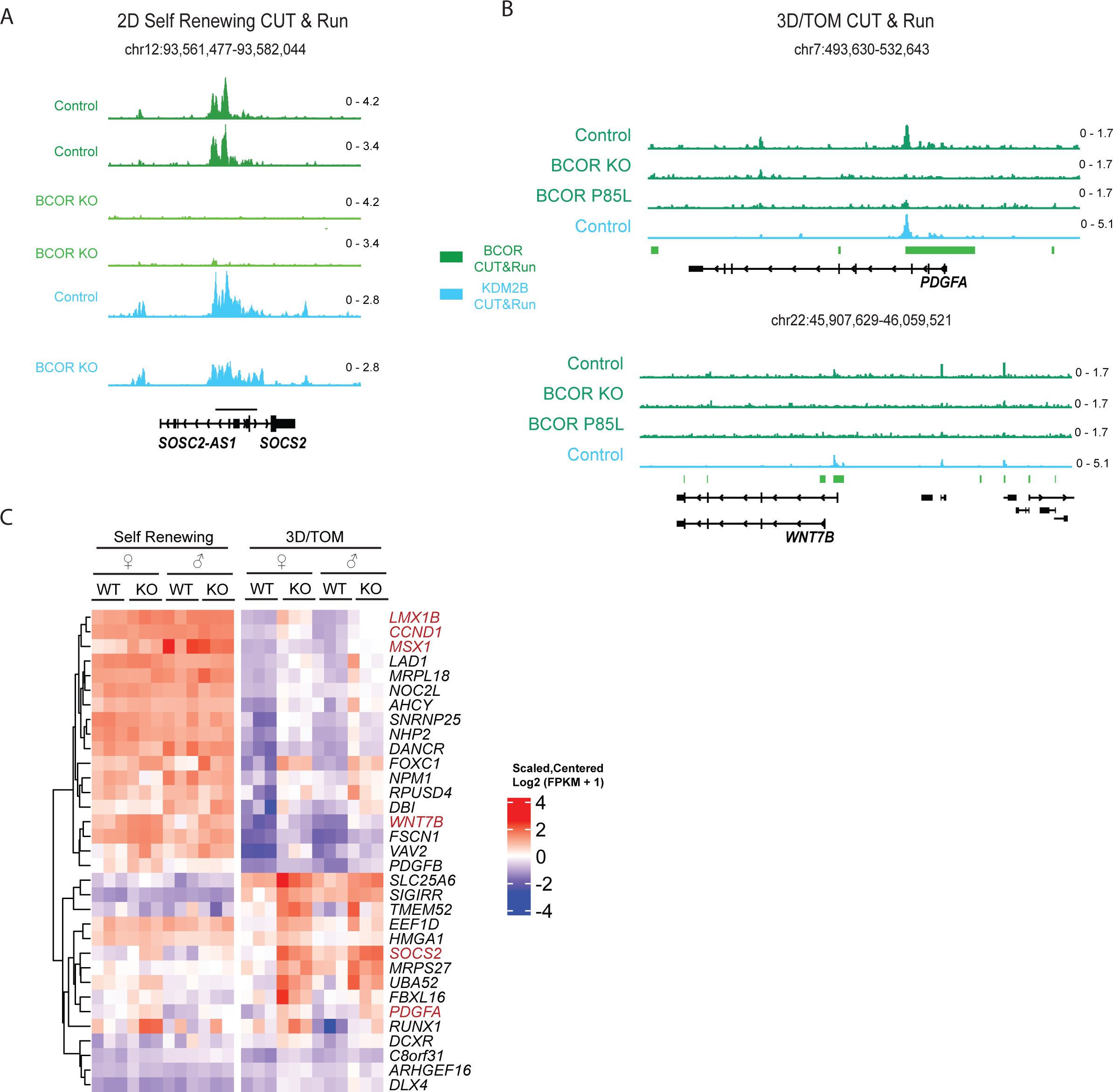
Genome Wide Occupancy of BCOR in human trophoblasts. A) CUT&RUN profiling of BCOR and KDM2B at the promoter of the *SOCS2* gene in control and *BCOR* knockout cell lines under self-renewing conditions. B) CUT&RUN of BCOR (green) and KDM2B (cyan) in 3D cell culture with TOM media. For BCOR, control, *BCOR* knockout and P85L missense mutant lines are shown. C) Expression heatmap of genes bound by BCOR in self-renewing or 3D/TOM cell culture conditions that also have upregulated gene expression, i.e. functional targets. Genes labeled in red were also functional targets in the mouse *in vivo*.

We then profiled BCOR occupancy of cells growing for four days in 3D culture conditions which showed a larger number of differentially expressed genes upon BCOR loss compared to 2D self-renewing cells. We observed 2,105 peaks present in control CT29 cells that were not detected in knock-out cell lines (**Fig. 4B**). As expected from the reduced abundance in western blots (**Fig. 3A**), the BCOR peak intensity in the P85L mutant CT29 cell line was greatly reduced (**Fig. 4B**) but still detectable at 375 of the 2,105 BCOR peaks. Combining peaks from cells cultured in self-renewing and differentiating in 3D/TOM media we identified 3,414 regions bound by BCOR in one or both conditions. Annotation revealed a similar enrichment for promoter regions as was observed in mouse ChIP seq (**Fig. 2E**). Among the genes bound by BCOR, we identified 33 genes that were upregulated under self-renewing or 3D/TOM differentiation. Hierarchical cluster analysis FPKM expression values from these genes identified two main patterns (**Fig. 4C**). Eighteen genes, including *LMX1B*, *WNT7B*, and *MSX1* were expressed in self-renewing conditions but were not completely repressed upon differentiation. In contrast, 15 genes including *PDGFA*, *SOCS2*, and *SLC26A6* were expressed at low levels in self-renewing cells and have higher expression in BCOR knock-out cells than control cells as they differentiate. Six of the 33 differentially expressed genes directly bound by BCOR were also bound and differentially expressed in mouse placenta (red labels, **Fig. 4C**) suggesting that functional targets of BCOR are also conserved between *in vivo* mouse models and cultured human trophoblasts.

We next sought to understand how cell fates are affected by BCOR loss in 3D cell culture using TOM media. We first performed immunofluorescence imaging of control and BCOR knockout cell lines after eight days of differentiation. Like our 2D directed experiments differentiating hTS cells into syncytiotrophoblast and extravillous trophoblast, we found that control and knockout cells were both capable of generating organoid-like structures with a layer of cytotrophoblast cells expressing CHD1 surrounding a core of syncytiotrophoblast cells expressing hCG (**Fig. 5A**). We did not detect a consistent difference between control and BCOR knockouts due to the highly variable shapes and sizes of the structures. To determine the number of cells of each type quantitative, we performed single cell profiling of control and knock-out cell lines. To profile syncytiotrophoblasts as single cells prior to cell-cell fusion events that would increase cell size and precluding their recovery in our cell isolation protocol, we profiled gene expression using the Parse split-seq platform on day four of differentiation in 3D culture using TOM media. We obtained high quality expression profiles for 2804 female controls, 1202 female BCOR knockouts, 389 male controls, 924 male BCOR knock-outs, and 1148 BCOR P85L missense male cell lines. After integration of the data across male and female cell lines we observed four statistically distinct clusters of cells (**Fig. 5B**). Three of these clusters expressed cytotrophoblast markers such as *TEAD4* and one expressed syncytiotrophoblast marker *CGA* (**Fig. 5C**). RNA velocity analysis shows cells moving slowly (smaller vectors) through the cytotrophoblast populations before quickly (larger vectors) transitioning to syncytiotrophoblast and stabilizing (smaller vectors). The proportion of cells within each cluster varied by parental line/sex and genotype (**Fig. 5D**). Notably, control male and female cell lines had a higher percentage of syncytiotrophoblast than *BCOR* knock-out lines. Together these results suggest that gene expression changes due to loss of *BCOR* have a modest effect on differentiation of trophoblast progenitor cells into syncytiotrophoblast cells

**Figure 5.**
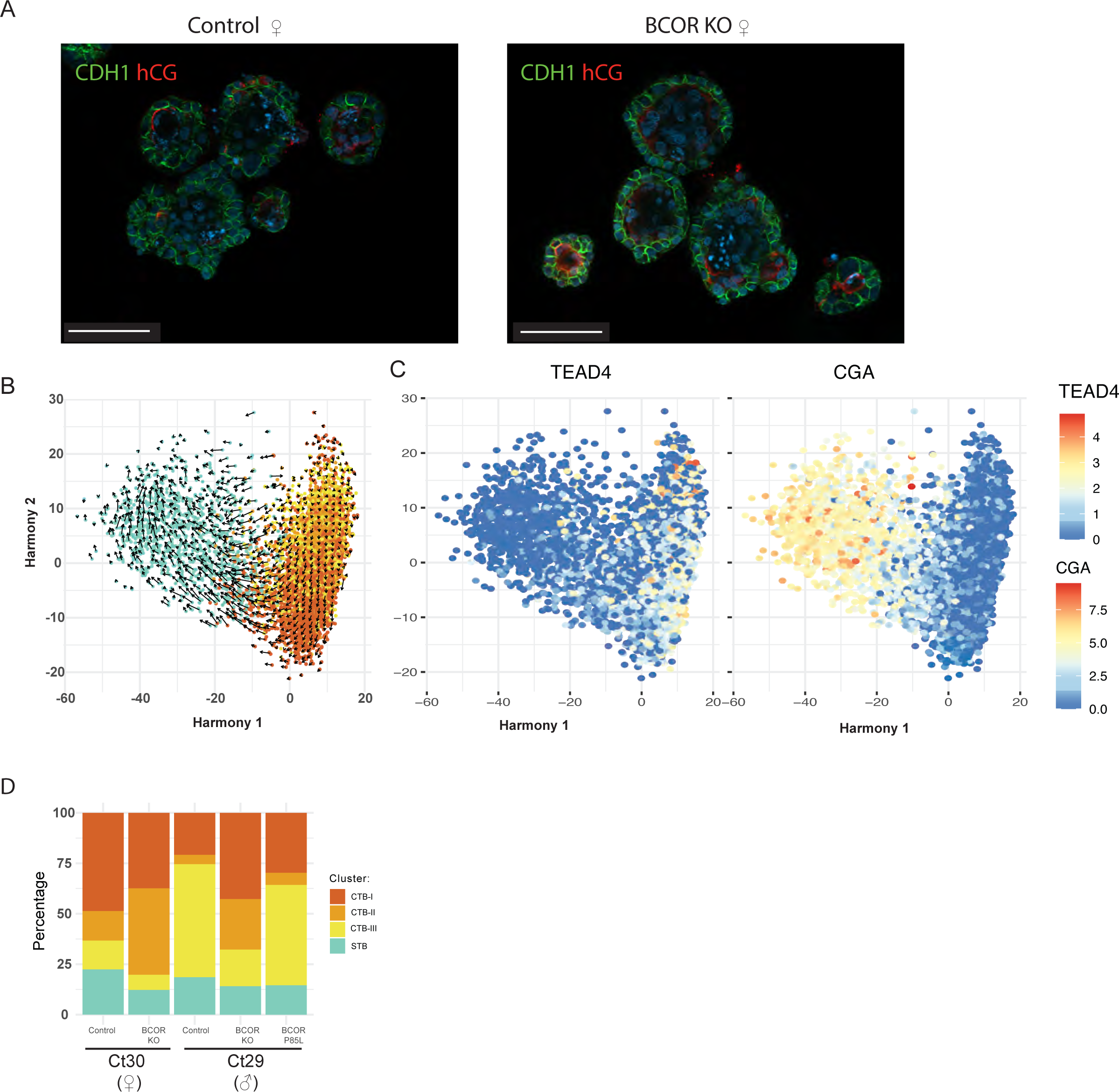
Single Cell Analysis of Autonomous Human Trophoblast Differentiation. A) CHD1 (green), hCG (red), and DAPI staining of control and BCOR knockout cell lines grown in 3D cell culture using TOM media. Trophoblast stem cells expressing CHD1 cells surround hCG expressing syncytiotrophoblast cells in organoid-like colonies grown after 8 days of differentiation. Scale bar is 100 μm. B) 2D projection of male and female datasets integrated with Harmony. RNA velocity (black arrows) shows cells differentiating from cytotrophoblast to syncytiotrophoblast. C) Cluster identities were confirmed by the expression of *TEAD4* and *CGA*. B) Relative proportions of cells in each cluster by genotype.

## Discussion

Gene expression profiling and genome-wide occupancy studies have identified a number of functional targets bound and repressed by BCOR in mice and human trophoblasts. Notably *WNT7B* and *PDGFA* have prominent BCOR peaks at their promoters are upregulated in both species likely impacting WNT and PDGF signaling pathways which are known to play an important role in placenta development(Hoch and Soriano 2003; Knöfler and Pollheimer 2013). BCOR also plays a role in silencing stem cell markers such as *Satb1* in mice and *HMGA2* in humans suggesting that loss of BCOR compromises the ability to differentiate appropriately. Lastly, BCOR expression in the placenta maintains lineage fidelity by repressing genes that promote alternate cell fates. *LMX1B* and *MSX1* are normally expressed at very low levels in control mice and human trophoblasts but are among the strongest upregulated genes in both species. These transcription factors are required for appropriate limb, kidney and craniofacial development(Chen et al. 1998; Satokata and Maas 1994). The promoters of both genes are bound by BCOR (**Fig. 4B,C**) although the increased abundance of WNT7B and faulty activin signaling due to BCOR loss (see below) may also play a role in aberrant expression of alternate lineage genes.

The fact that only 13 genes were differentially expressed in hTS cells under 2D self-renewing conditions suggests that BCOR is not required to maintain the stem cell population. This may have been fortuitous in that it allowed us to isolate clonal loss-of-function lines. Culturing control and *BCOR* knockout lines in 3D using TOM media revealed considerably more differentially expressed genes. Notably, this media contains 10-fold less of the TGFβ inhibitor, A83-01, than the 2D self-renewing media used for clonal selection (5 μm vs 0.5 μm). Lowering the inhibitor concentration should increase activin signaling and perhaps promote syncytiotrophoblast formation, analogous to what has been observed in mice (Natale et al. 2009). If BCOR repression is dependent on activin signaling in humans as previously shown in mice (X. Zhu et al. 2015), loss of BCOR may phenocopy impaired activin signaling to inhibit syncytiotrophoblast formation. Further experiments manipulating activin signaling pathways using 3D hTS cell culture models in the presence and absence of BCOR may prove useful in elucidating these mechanisms in human cells.

## Acknowledgments

This work was supported by awards from the National Institute for Child Health and Development: R01HD084459 to VJB and R21HD102770 to MDG.

## Methods

### Mouse Breeding & Genotyping

*Bcor* ^Flox910/Flox910^ female mice (Jackson #035713), backcrossed onto a C57BL/6 background, were bred to β-actin-Cre (Jackson #033984) males (Lewandoski, Meyers, and Martin 1997)(a gift of M. Lewandoski). All mice were maintained in conventional housing facilities. Presence of a copulation plug in the morning was recorded as day E0.5. All experimental protocols involving mice described in this publication have been approved by the University of Minnesota Institutional Animal Care and Use Committee.

### RNA Sequencing

Mouse RNA-seq was performed on individual placentas as was done previously in mouse T cells(Kotov et al. 2019). Embryos were used for genotyping with PCR primer sequences described previously(Hamline et al. 2020). Extraembryonic mouse tissue (E6.5-E9.5) was dissected away from embryonic tissue. hTS cells were grown per their experimental condition and removed from their growth medium. All RNA was extracted and separated using TriZOL (Ambion) reagent and RNA purification columns (Qiagen). Libraries were prepared with the KAPA mRNA hyperprep kit (Roche). Samples were then gel purified and size selected using E-gel EX 2% Agarose (Invitrogen) and was followed by gel extraction (Qiagen). Once libraries passed internal quality control they were sequenced on the Illumina platform.

### Paraffin-embedding, H&E, and Immunofluorescence

Tissues were fixed between 15 minutes to 3 hours at 4°C in 4% paraformaldehyde (PFA, Electron Microscopy Sciences). Post-fixation samples were dehydrated with ethanol, cleared with Citrisolv (Decon Labs), and infiltrated with Paraplast Xtra (Leica Biosystems) paraffin wax. Once embedded samples were sectioned between 5-7µm using a Reichert-Jung Microtome.

Sections embedded in paraffin were deparaffinized with Cirtisolv and rehydrated. H&E staining was performed with Gills Type 2 Hematoxylin (Sigma) and 1:1 Eosin (Fisher Scientific): 80% ethanol. For tissue used for immunofluorescence, a Citric Acid Buffer antigen retrieval step was optimized for specific primary antibodies ranging from 20 minutes to 1 hour. Sections were then permeabilized using 0.1% Tween20 in PBS. Sections were stained with primary antibodies, 1:400 Anti-Monocarboxylate Transporter 1 (MCT1, Sigma AB1286-I), and 1:400 Trophoblast specific protein alpha (TPBPα, Abcam ab104401) overnight at room temperature. Slides were washed thoroughly with PBS before incubating with the secondary antibodies, donkey anti-rabbit 488 (Biotuim), and donkey anti-chicken 568 (Invitrogen). All antibody incubations were done in block solution containing 5-10% Normal Donkey serum and 3% Bovine Serum Albumin (BSA, Sigma). After early initial staining, it was realized that some extraembryonic tissue was autofluorescent. Subsequent slides were incubated with Trueblack Lipofuscin Autofluorescence Quencher (Biotium). Images were acquired on an AxioImager (Zeiss).

### Cryo-embedding and Immunofluorescence

Tissues were fixed with 4% PFA between 1 hour at room temperature and overnight at 4C. They were then washed with PBS, and equilibrated in 30% sucrose to cryo-protect samples. Cryo-protected samples were placed in O.C.T. Compound (O.C.T., Tissue-Tek) and snap-frozen using dry ice. Once embedded samples were stored at -80°C until sectioned between 7-15µm on a Bright Instrument Company cryostat. PBS was used to wash O.C.T. from sections which were then permeabilized with 0.2% Triton X-100. Primary staining, in-house BCOR, was done overnight at 4°C. Slides were washed thoroughly with PBS before incubating with the secondary antibodies, Donkey anti-rabbit 488 (Invitrogen). All antibody incubations were done in block solution containing 5-10% Normal Donkey serum and 3% BSA. Images were acquired on an AxioImager (Zeiss).

### Culture of 2D hTSCs

Two patient-derived lines of Human Trophoblast Stem Cells (hTSCs) were used in this study, CT29, and CT30 (gifts from H. Okae), and were cultured using a modified version of the published protocol (Okae et al., 2018). hTSCs were maintained in a self-renewing condition on plates coated in iMatrix511 (Matrixome) grown in a DMEM/F12 (Gibco) TS Complete media containing 100 µg/mL Primocin (InvivoGen), 0.15% BSA (Sigma), 1% Insulin-Transferrin-Selenium-Ethanolamine 100x (ITS-X, Gibco), 1% Knockout Serum Replacement (KSR, Gibco), 0.2 mM L-ascorbic acid (Sigma), 2.5 µM Y27632 (R&D/Tocris), 25 ng/mL rhEGF (R&D/Tocris), 0.8 mM Valproic Acid (Amsbio), 5 µM A83-01 (Tocris Bioscience), and 2 µM CHIR99021 (Tocris Bioscience). Cells were incubated at 37°C in 5% CO_2_ and 3-20% O_2_. Cells were passaged using TrypLE Express Enzyme (Gibco), and stored in long and short-term storage in Cellbanker 1 (Zenoaq).

### Generation of hTSC BCOR knockouts and P85L point mutant

Three unique guide RNAs were designed to target *BCOR* in both CT29s and CT30s in addition to three control guides including two guides targeting non-coding regions of the X-chromosome and a GFP sequence not present in the genome.. These guides were introduced to the cells via lentivirus transduction. Once the cells were selected for guide incorporation using a puromycin-selectable marker, they were dissociated using TrypLE and resuspended in a Sucrose/MgCl_2_ buffer at 2k cells/µl. 50k cells were removed from the Sucrose/MgCl_2_ and added to 10 µl of Resuspension buffer for electroporation. 200 ng CleanCap Cas9 (5moU) mRNA (Trilink, L-7206) was electroporated into the cells using the Neon™ Transfection System (ThermoFisher). CleanCap eGFP mRNA was used as a control. Cells were then allowed to recover without antibiotics, Primocin, or puromycin until a healthy pool was established. The pools were then diluted out to single colonies on a 96-well plate, and colonies were picked and expanded if they appeared healthy. All clones were incubated in TS Complete media and at 37°C in 5% CO_2_ and 3-20% O_2_. The BCOR knockout clones were then verified by Western blot and Sanger sequencing.

### Directed hTSC syncytiotrophoblast (STB) differentiation

Self-renewing cells were seeded at double the normal seeding concentration with TS complete media, and iMatrix511, and were incubated overnight on Day 0. On Day 1 the media was changed to TS STB Differentiation media containing DMEM/F12 with 100 µg/mL Primocin, 0.1% BSA, 1% ITS-X, 0.1mM 2-Mercaptoethanol (BME, Gibco), 2.5 µM Y27632, 4% KSR, and 2 µM forskolin (Sigma). STBs were incubated at 37°C in 5% CO_2_ and 3% O_2_. Differentiation was confirmed with an at-home HCG test (sensitivity: 25 mIU/mL)(Pregmate), and harvested on Day 4.

### Directed hTSC extravillous trophoblast (EVT) differentiation

Wells were first coated in 1 ug/mL Col IV for 1.5 hours. Self-renewing cells were seeded at normal seeding concentrations on Day 0 with TS EVT differentiation media including DMEM/F12 with 100 ug/mL Primocin, 0.1% BSA, 1% ITS-X, 0.1mM BMEl, 2.5 µM Y27632, 4% KSR, 7.5 µM A83-01, 0.1µg/mL NRG1-beta1 (Cell Signaling Technology). Matrigel (Corning) 20µl/mL was added on Day 0. Media was changed to TS EVT differentiation media without NRG1-beta1 on Day 3 and matrigel 5 µl/mL was added. On Day 6 media was changed to TS EVT differentiation media without NRG1-beta1 and KSR. Matrigel 5 µl/mL was added. EVTs were incubated at 37°C in 5% CO_2_ and 3-20% O_2_. Cells were harvested either on Day 4 or Day 8.

### hTSC 3D /TOM Differentiation

Differentiation of hTSCs in 3D cell culture using TOM media was achieved by seeding self-renewing hTSCs in Matrigel droplets. Seeding density was set at 8,000 cells per 30 ul droplet of 78% Matrigel. Five droplets were added to a 12-well plate and covered with 1 mL of TOM [Advanced DMEM/F12 (Gibco), 1x N2 supplement (Gicbo), 1x B27 supplement minus vitamin A (Gibco), Primocin 100 μg/mL, *N*-Acetyl-L-cysteine 1.25 mM (Sigma), L-glutamine 2 mM (ThermoFisher), recombinant human EGF 50 ng/mL, CHIR99021 1.5 µM, recombinant human R-spondin-1 80 ng/mL (Peprotech), recombinant human FGF-2 100 ng/mL (Peprotech), recombinant human HGF 50 ng/mL (Peprotech), A83-01 500 nM, prostaglandin E_2_ 2.5 µM (Sigma), Y-27632 2 µM.] Organoids were incubated at 37°C in 5% CO_2_ and 3% O_2_. hTS cells were seeded on day 0, media was changed on day 2, and harvested for analysis on day 4.

### 3D/TOM Immunofluorescence

Organoids used for imaging were seeded with 3,000 hTS cells in one 30 ul droplet of 78% Matrigel on an 8-well Ibidi treat coverslip (Ibidi) and fed with TOM on days 2, 4, 6, and 7. On day 8 the cells were fixed for 10 minutes with 4% PFA, permeabilized with 0.3% Triton-x100, and stained with 1:200 E-cadherin (BD Transduction Laboratories) and 1:200 HCG (Abcam) overnight in 10% donkey block (10% normal donkey serum, 3% BSA and 0.1% Tween20). Organoids were incubated with secondaries, donkey anti-mouse 488 (Invitrogen), and donkey anti-rabbit 594 (Invitrogen) in 10% donkey block for 2 hours and DAPI for 15 minutes. Organoids were then stored at 4°C in PBS until imaging on an AxioObserver using the Apotome 3 (Zeiss) for optical sectioning.

### Single Cell RNA Sequencing of cells in 3D/TOM media

Differentiated organoid-like cells were removed from Matrigel using Cell Recovery Solution (Corning). Once free from Matrigel the cells were digested into a single-cell suspension using TrypLE and DNAse I (Worthington Biochemical Corporation). Cells were filtered through a 20% Percoll (Sigma) layer followed by a 100um filter and a 40um Flowmi® filter. Single cells were only kept if the viability was greater than 75% via AO/PI staining. Cells were fixed and frozen using the Parse Biosciences Cell Fixation kit. Transcripts were barcoded and libraries were made using the Parse Biosciences Evercode Mini.

### ChIP-seq and CUT&RUN

ChIP-seq in mouse placenta was performed as was done previously in mouse T cells(Kotov et al. 2019). CUT&RUN was performed using a modified CUT&RUN protocol. Cells were grown per condition and single cells were isolated. About 100k cells per condition were bound to Concanavalin A beads (Bangs Laboratories). Cells were permeabilized using Digitonin. Primary antibodies, BCOR (AB_2716801) and KDM2B(AB_2716799) were incubated for 2 hours at room temperature on a shaking platform. Cells were then incubated with an in-house pA/G-MNase for 1 hour on ice. CaCl_2_ was added to activate the pA/G-MNase. CUT&RUN fragments were released and incubated overnight at 55°C with Proteinase K and SDS. The following day the CUT&RUN fragmented DNA was extracted with Phenol Chloroform. Libraries from the CUT&RUN fragments were made with a KAPA Hyper Prep kit (Roche). Samples were then gel purified and size selected using E-gel EX 2% Agarose (Invitrogen) followed by gel extraction (Qiagen).

### Western Blot

2D self-renewing hTSCs were washed with PBS and harvested in TriZOL. Proteins were prepped and resuspended in NuPAGE LDS buffer (Invitrogen) and BME. Samples were run on NuPAGE 3-8% Tris Acetate Gel (Invitrogen) in Tris-Acetate SDS Running Buffer (Novex). Blots were transferred overnight at 4°C using NuPAGE Transfer Buffer (Novex) and methanol. Blocking was performed with Intercept Blocking Buffer TBS (LI-COR). Blots were stained overnight at 4°C with our in-house BCOR antibody and a β-actin (Santa Cruz) antibody in the Intercept block. Secondaries were incubated for 1-2 hours at room temperature. Blots were scanned on an Odyssey M (LI-COR).

